# FtsH Protease Inactivation Allows Accumulation of Aberrant Photosystem II in a *Chlamydomonas* Rubredoxin Mutant

**DOI:** 10.1101/2022.02.24.481860

**Authors:** Robert H. Calderon, Catherine de Vitry, Francis-André Wollman, Krishna K. Niyogi

## Abstract

The assembly of photosystem II (PSII) requires the participation of assembly proteins that facilitate the step-wise association of its protein and pigment components into a functional complex capable of oxidizing water and reducing plastoquinone. We previously identified one such factor, the membrane-bound rubredoxin RBD1, but its precise role remains unknown in part due to the inability of the *2pac* mutant strain of *Chlamydomonas reinhardtii*, which lacks RBD1, to accumulate PSII. Here, we show that decreased PSII accumulation in *2pac* is due to increased proteolytic degradation. Inactivating the thylakoid membrane FtsH protease in the *2pac* mutant background led to an increase in the abundance of PSII subunits and their integration into higher molecular weight complexes, including PSII dimers, capable of sustaining photoautotrophic growth. Dark- and low light-grown *2pac ftsh1-1* both accumulated a 23-kD fragment of the D1 protein, a marker typically associated with structural changes resulting from photodamage or photoinhibition. We introduced a HIS-tagged version of the PsbH protein into the *2pac ftsh1-1* background to purify and examine PSII. We found no detectable changes with respect to cofactor composition relative to the wild-type, leading to us to propose a model in which RBD1 promotes the proper folding of D1, possibly via delivery or reduction of the non-heme iron during PSII assembly. Our results demonstrate that introduction of the *ftsh1-1* mutation into mutants defective in the biogenesis of thylakoid membrane complexes can allow for the accumulation and study of aberrant complexes that would otherwise be degraded due to their high protease sensitivity.

## Introduction

Photosystem II (PSII) is a light-driven water:plastoquinone oxidoreductase essential for the growth of all oxygenic photoautotrophic organisms. In generating oxygen as a byproduct, it also enables the growth of most, if not all, aerobic organisms on the planet. The assembly of PSII occurs in a step-wise process that requires a variety of assembly factors, some of which are conserved from cyanobacteria to higher plants (de Vitry et al., 1989; Nixon et al., 2010; Komenda et al., 2012; Nickelsen and Rengstl, 2013; Spaniol et al., 2022). In a previous publication, we isolated and characterized cyanobacterial, algal and plant mutants lacking one such assembly factor, a highly conserved rubredoxin called RubA in cyanobacteria or RBD1 in photosynthetic eukaryotes (Calderon et al., 2013). Rubredoxins are small iron-containing proteins that function in electron transfer reactions. The RubA/RBD1 protein is associated with thylakoid membranes (Shen et al., 2002) and required for the assembly of PSII in *Synechocystis sp*. strain PCC 6803, *Chlamydomonas reinhardtii*, and *Arabidopsis thaliana* (Calderon et al., 2013), and it was later determined to be associated with intermediate complexes during PSII assembly rather than mature, fully assembled PSII (García-Cerdán et al., 2019; Kiss et al., 2019). In *Chlamydomonas*, RBD1 appears to have roles both in assembly of PSII intermediate complexes and in photoprotection during PSII assembly and repair (García-Cerdán et al., 2019). However, further elucidation of the specific role of RBD1 has been limited by our inability to isolate and study PSII that assembles in the absence of RBD1 because of the low accumulation of PSII subunits and complexes in *rbd1* mutants.

The FtsH protease complex is a membrane-localized ATP-dependent protease found in all oxygenic photoautotrophs (Lindahl et al., 1996). It has a well-characterized role in the degradation of damaged PSII subunits during photoinhibitory and oxidative stresses (Lindahl et al., 2000), and it has also been implicated in the degradation and remodeling of cytochrome *b*_6_*f* (cyt *b*_6_*f*) complexes (Malnoë et al., 2014). In *Chlamydomonas*, the thylakoid membrane-anchored FtsH protease exists as a heterocomplex comprised of FtsH1 and FtsH2 isoforms (Malnoë et al., 2014). Strikingly, two *Chlamydomonas ccb* mutants, which normally lack cyt *b*_6_*f* complexes due to mutations in cyt *b*_6_*f* assembly factors, were found to be able to accumulate aberrant forms of cyt *b*_6_*f* in a *ftsh1*-mutant background (Malnoë et al., 2011).

Here we have examined the role of the RBD1 protein in the assembly of PSII in *Chlamydomonas* by inactivating the FtsH protease in the *2pac* mutant, which lacks RBD1. We show that combining the *ftsh1-1* and *2pac* mutations permits the accumulation of a misassembled variant of PSII. We find that accumulation of this variant PSII is sufficient to allow photoautotrophic growth in the absence of RBD1, and thus we show that inactivation of FtsH can be a powerful tool for studying the role of PSII assembly factors.

## Results

### The lack of PSII in *2pac* is due to instability of PSII subunits rather than a defect in translation

Given that the assembly of PSII monomers occurs to a much lower extent in the *2pac* mutant than in the wild type (García-Cerdán et al., 2019), we sought to investigate whether the decrease in PSII accumulation might be caused by a defect in translation of chloroplast-encoded PSII subunits. We therefore pulse-labeled *Chlamydomonas* cells for several minutes in order to detect rates of protein translation rather than rates of degradation. Wild-type (4A+ or T222), *2pac*, and the Δ*psbA* strain Fud7 (Bennoun et al., 1986) cells were grown on acetate as a carbon source and allowed to reach logarithmic growth phase before being pulse-labeled with [^14^C]-acetate for 7 min in the presence of cycloheximide (to inhibit cytosolic translation). As shown in **Fig. 1A**, there were no major differences in the incorporation of the radioactive signal into newly synthesized proteins between 4A+, T222, *2pac*, and Fud7 (with the exception of D1 for Fud7). The assignment of the PSII proteins D1 (PsbA), D2 (PsbD), CP43 (PsbC) and CP47 (PsbB) is based on mutant analysis (de Vitry et al., 1989). Levels of radiolabeled CP47, CP43 and cytochrome *f* are slightly lower in *2pac* relative to both WT strains and Fud7, but these proteins are clearly being actively translated.

**Figure 1:**
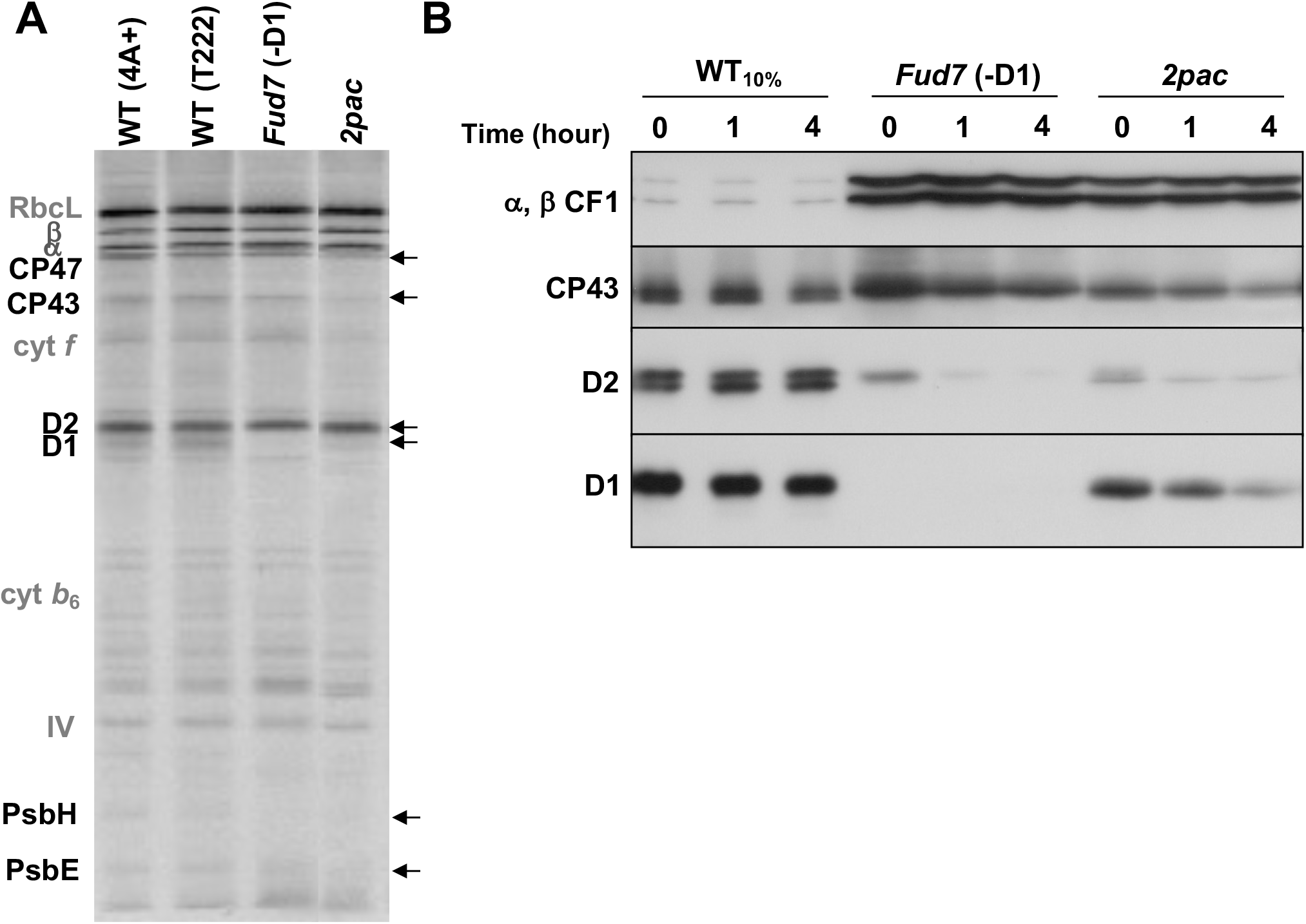
PSII subunits are translated, but unstable in the absence of RBD1. **A)** Plastid-encoded proteins from [^14^C]-acetate-radiolabeled wild-type (4A+ and T222), Fud7 (Δ*psbA*), and *2pac* whole-cell extracts separated by SDS-PAGE on a 12-18% polyacrylamide gel in the presence of 8 M urea and visualized by autoradiography. PSII subunits are in black and indicated by arrows. Other photosynthetic proteins are labeled in gray. **B)** Immunoblot analysis of steady-state levels of α and β subunits of plastid ATP synthase (α, β CF1), D1, D2, and CP43 after 0, 1, or 4 hours of incubation with chloroplast translation inhibitors. All loaded samples contained 20 μg chlorophyll, with the exception of the “WT_10%_”, which contained 2 μg chlorophyll.

Because subunits of PSII are being translated in *2pac*, we hypothesized that the lack of accumulation of PSII complexes might be due to instability of PSII subunits. To test this, wild-type, Fud7, and *2pac* cells were allowed to reach logarithmic growth phase before adding chloroplast translation inhibitors (lincomycin and chloramphenicol). Samples were collected at three time points (0, 1, and 4 h after incubation) for immunoblot analysis. These immunochase data showed that, over the time course, the PSII subunits D1, D2, and CP43 all rapidly disappeared in *2pac*, whereas they remained stable in wild-type cells (**Fig. 1B**). In the Fud7 strain, D1 was absent, and D2 and CP43 were unstable, as expected (de Vitry et al., 1989). In contrast to PSII proteins, the α and β subunits of the chloroplast ATP synthase were stable in all three strains (**Fig. 1B**). The data therefore suggest that genes encoding thylakoid membrane proteins are all transcribed and translated in *2pac*, and the lack of PSII is due to the specific degradation or instability of PSII subunits relative to subunits of other photosynthetic complexes.

### *2pac ftsh1-1* accumulates higher levels of PSII subunits and complexes than *2pac*

Given the well-established role of FtsH in the degradation of damaged PSII, we hypothesized that the introduction of the *ftsh1-1* mutation into the *2pac* strain could result in a decrease in the degradation of PSII subunits and thus an increase in steady-state levels of these proteins. To test this, we first crossed the *2pac* and *ftsh1-1* strains and isolated *2pac ftsh1-1* double mutants (**Supplemental Fig. 1**). We then measured the accumulation of PSII subunits in these strains by immunoblot analysis after growth in low light. As shown in **Fig. 2**, there was a marked increase in the amounts of D1, D2, CP43, and CP47 in the *2pac ftsh1-1* strain relative to both the parental *2pac* strain and the *2pac* progeny isolated from the same tetrad, although the levels of these PSII subunits were not as high as in the wild-type or *ftsh1-1* strains. This pattern was observed in all tested *2pac ftsh1-1* progeny **(Supplemental Fig 2)**. In addition, immunodetection with an antibody raised against the loop connecting helices D and E of the D1 protein (α-DE loop) revealed a band at 23 kD in *2pac ftsh1-1* and *ftsh1-1* (**Fig. 2**). This fragment has been detected in the single *ftsh1-1* mutant only in the light and under various stress conditions (Malnoë et al., 2014). It has been attributed previously to a photodamage-induced cleavage product of the full-length D1 protein (De Las Rivas et al., 1992) as well as a translational pause intermediate of D1 (Mullet et al., 1990). In contrast, the wild-type strains did not accumulate any of the 23-kD fragment under the tested conditions. The intensity of the 23-kD fragment band (relative to full-length D1) was much higher in *2pac ftsh1-1* than in *ftsh1-1*, suggesting an increased PSII light sensitivity of *2pac ftsh1-1* compared to *ftsh1-1*.

**Figure 2:**
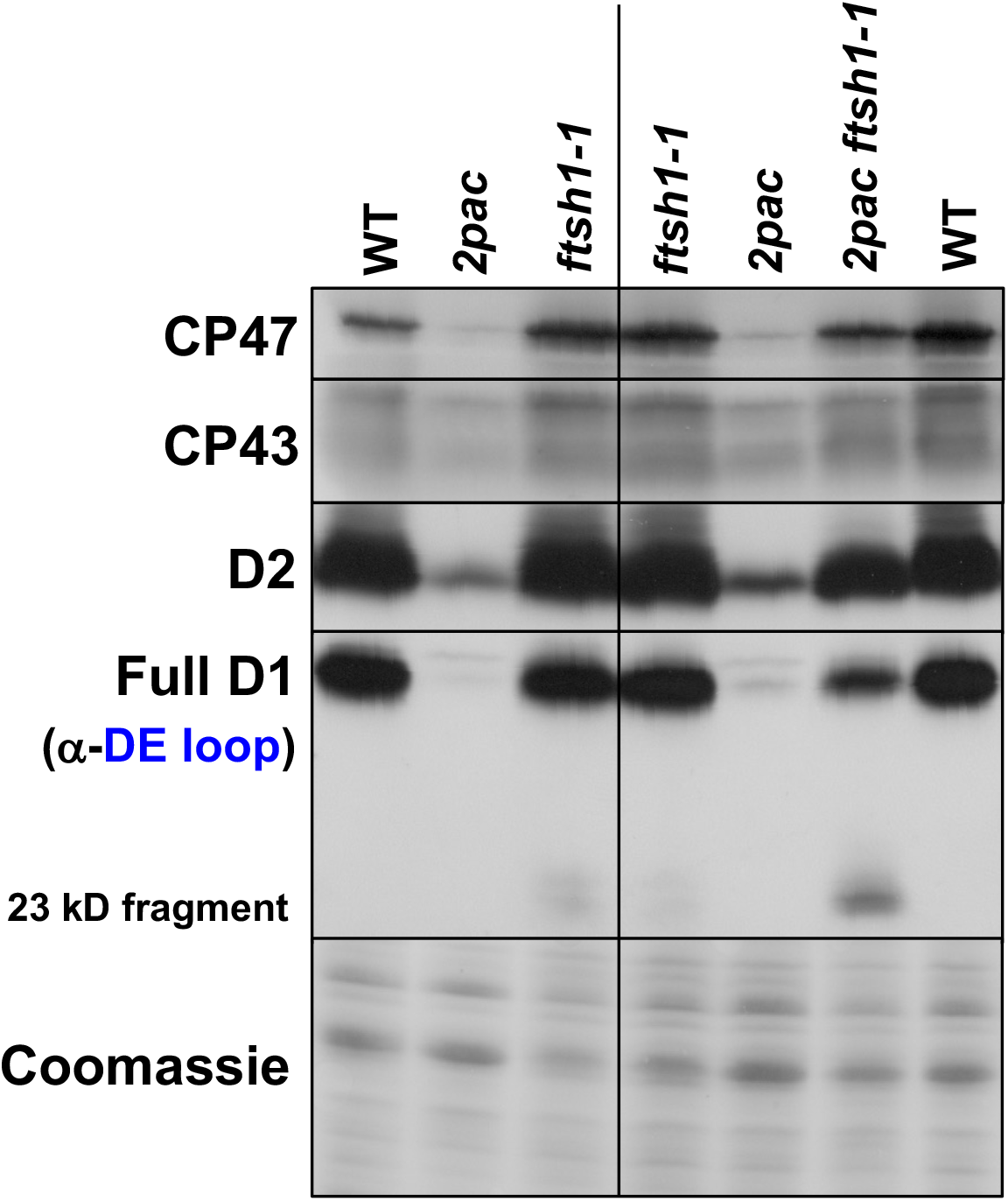
Inactivation of FtsH increases accumulation of PSII subunits in *2pac*. Immunoblot analysis of PSII subunit levels from a representative tetrad (tetratype tetrad 1, right side) obtained by crossing *2pac* and *ftsh1-1*, with parental strains and wild-type as controls (left side). Whole-cell protein extracts from the strains grown in acetate-containing TAP medium at 6 μmol photons m^−2^ s^−1^ were separated by SDS-PAGE on a 12-18% polyacrylamide gel in the presence of 8 M urea and immunodetected with antibodies against PSII subunits (CP47, CP43 and D2) and a peptide of the loop between the transmembrane helices D and E of D1 (D1-DE loop). Coomassie protein staining is provided as loading control.

To determine the assembly status of the PSII subunits in the *2pac ftsh1-1* mutant, we isolated thylakoid membranes from cells grown photoheterotrophically at 6 μmol photons m^−2^ s^−1^ and examined whether or not PSII subunits were being incorporated into complexes by 2D-PAGE analysis. In previous work with the *2pac* mutant, PSII subunits were found to assemble into PSII monomers but not dimers or supercomplexes (García-Cerdán et al., 2019). However, in the *2pac ftsh1-1* mutant, we detected these subunits in both PSII monomers and dimers (**Fig. 3**), indicating that these subunits were not only accumulating to greater levels **(Fig. 2)** but also being incorporated into higher molecular weight complexes. Additionally, in the *2pac ftsh1-1* mutant, some of the D2 subunit appeared to accumulate into a subcomplex of smaller molecular weight than the CP43-less PSII assembly intermediate RC47 complex (indicated by an arrow in **Fig. 3**). These smaller subcomplexes were not detected in the *ftsh1-1* mutant.

**Figure 3:**
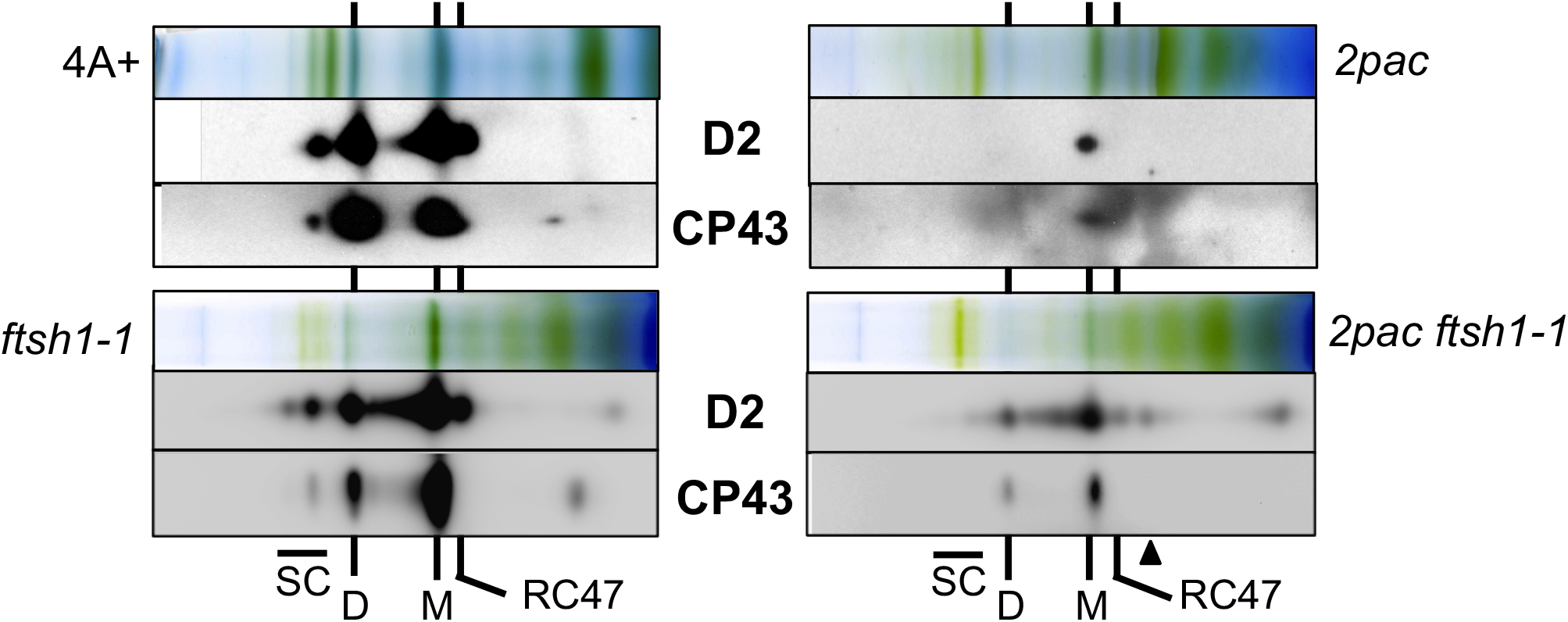
Two-dimensional blue native (BN)/SDS-PAGE analysis of PSII in *2pac ftsh1-1* relative to parental strains. Thylakoid membrane complexes from WT (4A+), *ftsh1-1, 2pac* and *2pac ftsh1-1* were solubilized with 1% β-DM and separated by BN-PAGE (top panels), followed by SDS-PAGE in the second dimension and immunoblot analysis to detect D2 and CP43 (lower panels). SC, PSII super complex; D, PSII dimer; M, PSII monomer; RC47, CP43-less PSII assembly intermediate (Komenda et al., 2004). Smaller complexes specifically detected in *2pac ftsh1-1* with antibodies against D2 are indicated by an arrow.

### *2pac ftsh1-1* grows photoautotrophically in low light and exhibits variable fluorescence

We next examined whether the PSII complexes that accumulate in *2pac ftsh1-1* can sustain photoautotrophic growth. As shown in **Fig. 4A**, the *2pac ftsh1-1* mutant was able to grow slowly under low light conditions on minimal medium, indicating that the PSII in this strain is at least partially functional. To test the activity of PSII under the same light conditions, we measured variable chlorophyll fluorescence to calculate the maximum efficiency of PSII (F_v_/F_m_) and found that *2pac ftsh1-1* exhibited low levels of variable fluorescence (**Fig. 4B**). The *2pac ftsh1-1* mutants displayed more variable fluorescence when grown at very low light on solid medium (**Fig. 4B** and **Supplemental Fig. 3**) than when grown at low light in strongly aerated liquid cultures (**Fig. 4C** and **Supplemental Fig. 4**), suggesting oxygen- and/or light-dependent loss of PSII activity.

**Figure 4:**
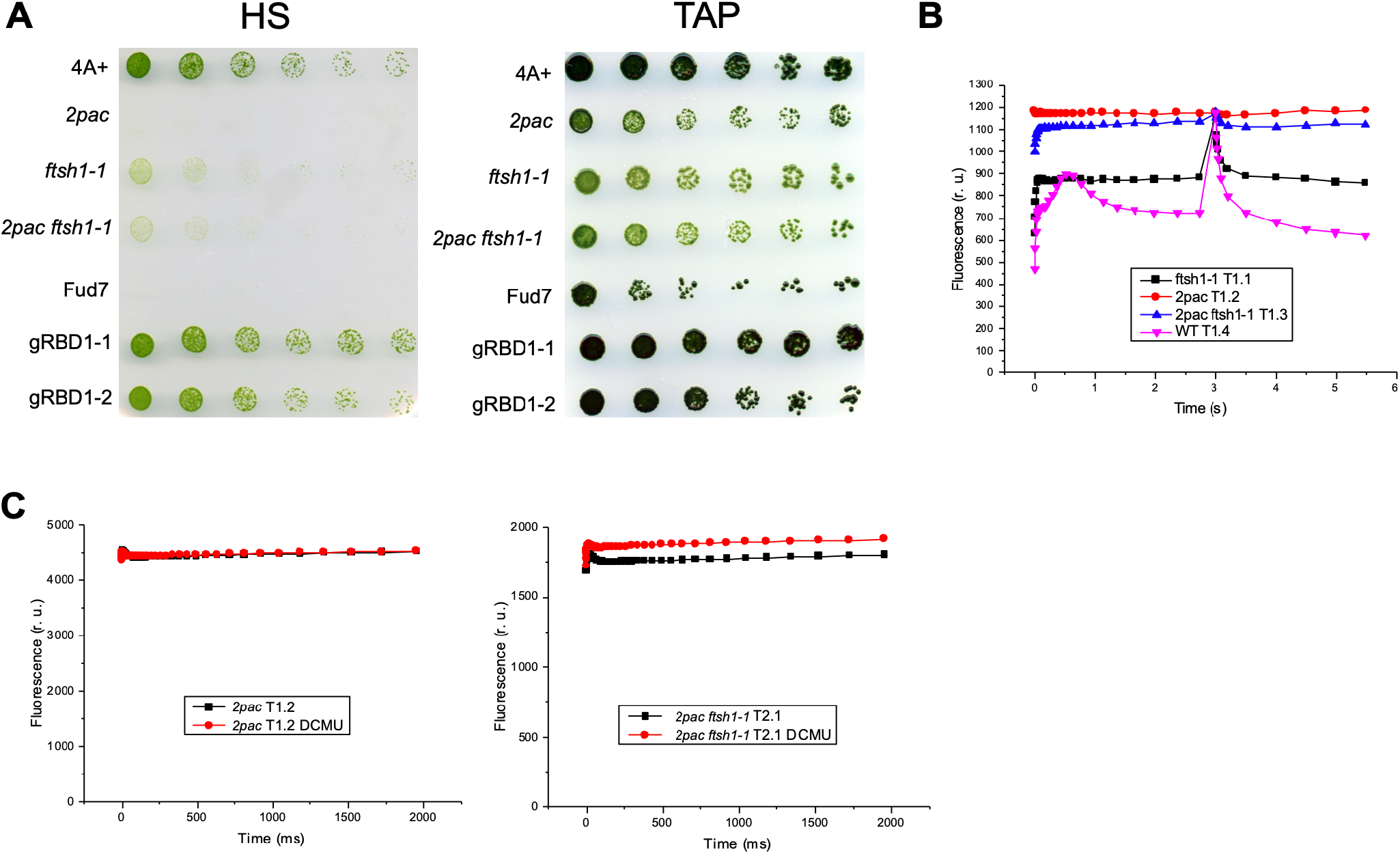
*2pac ftsh1-1* grows photoautotrophically and displays low variable fluorescence. **A)** Growth on plates without (HS, *left panel*) and with (TAP, *right panel*) acetate at 30 μmol photons m^−2^ s^−1^ of wild-type (4A+), *2pac, ftsh1-1, 2pac ftsh1-1*, a strain lacking the *psbA* gene encoding the D1 protein (Fud7), and two complemented lines (gRBD1–1 and gRBD1–2). **B)** Fluorescence induction kinetics in relative units (r. u.) at low actinic light of dark-adapted cells followed by a saturating pulse after 3 s to determine F_m_. Strains as in Fig. 2 from tetratype tetrad 1 grown on TAP plates at 6 μmol m^−2^ s^−1^. **C)** Fluorescence induction kinetics at low actinic light of dark-adapted cells in the presence (red) or absence (black) of the PSII-specific inhibitor DCMU. *2pac* and *2pac ftsh1-1* strains were grown in strongly aerated TAP liquid culture at 6 μmol m^−2^ s^−1^.

### Dark-grown *2pac ftsh1-1* displays variable fluorescence and accumulates a distinctive fragment of the D1 protein

We took advantage of the ability of *Chlamydomonas* to synthesize chlorophyll in the dark as well as in the light (Beale, 2009) to determine whether the PSII in *2pac ftsh1-1* might be light sensitive. To address this possibility, we grew the *2pac ftsh1-1* mutant in darkness and again assayed F_v_/F_m_ **(Fig. 5A**). The dark-grown mutant cells displayed an appreciable increase in F_v_/F_m_, consistent with higher levels of functional PSII.

**Figure 5:**
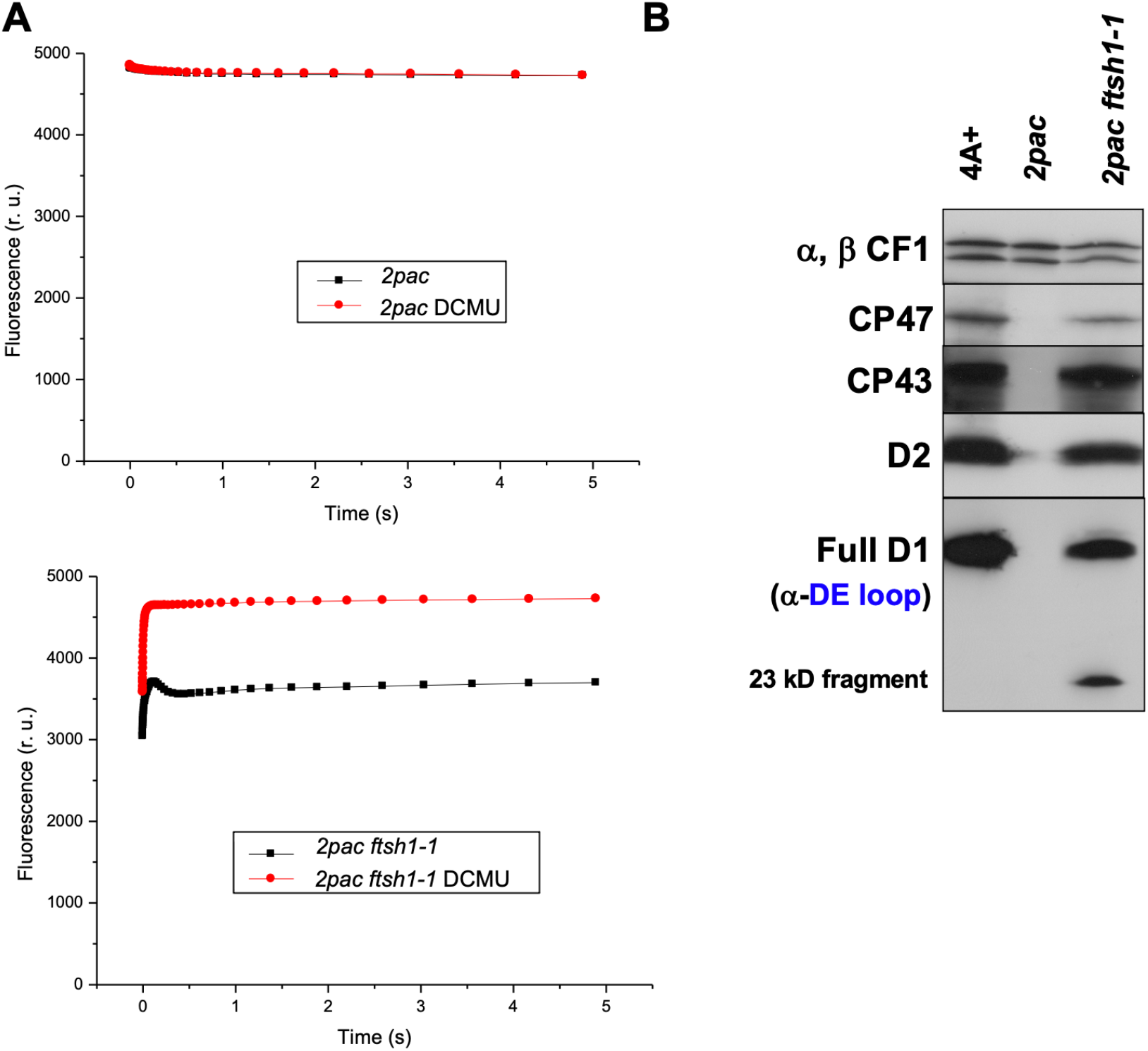
Dark-grown *2pac ftsh1-1* displays variable fluorescence and accumulates a 23-kD fragment of the D1 protein. **A)** Fluorescence induction curves of dark-grown liquid cultures of *2pac* (*upper panel)* and *2pac ftsh1-1* (*lower panel*) in the presence (red) or absence (black) of the PSII-specific inhibitor DCMU. An absence of DCMU-altered kinetics in *2pac* indicates an absence of variable fluorescence and PSII activity (F_v_/F_m_), whereas DCMU treatment reveals measurable F_v_/F_m_ in *2pac ftsh1-1*. **B)** Immunoblot analysis of PSII subunit levels from dark-grown WT (4A+), *2pac*, and *2pac ftsh1-1*. An antibody raised against a peptide from the stromal loop connecting helices D and E of the D1 protein (α-DE loop) recognizes both full-length D1 and a 23-kD N-terminal fragment, both of which accumulate in dark-grown *2pac ftsh1-1*.

We hypothesized that the low-light inactivation of PSII in *2pac ftsh1-1* might be due to either decreased abundance of PSII in the light or to a structural or conformational difference between these reaction centers and those from the wild-type. To test this, we first compared the accumulation of PSII subunits in dark-grown wild-type, *2pac*, and *2pac ftsh1-1* cells. As shown in **Fig. 5B**, the accumulation of PSII subunits is comparable between dark-grown wild-type and *2pac ftsh1-1* cells. While overall accumulation of mature proteins was comparable, we noted that immunodetection with the α-DE loop antibody revealed a band at 23 kD in dark-grown *2pac ftsh1-1* and not in the wild type **(Fig. 5B**), as was also the case in light-grown *2pac ftsh1-1* **(Fig. 2**). Because this D1 fragment is absent in the dark-grown single *ftsh1-1* mutant (Malnoë et al., 2014) and dark-grown *2pac* (**Fig. 5B**) but present in the dark-grown double *2pac ftsh1-1* mutant (**Fig. 5B**), it arises as a consequence of combining the *2pac* mutation with inactivation of the thylakoid protease FtsH.

### Purification and characterization of PSII from dark-grown *2pac ftsh1-1* cells

We hypothesized that the presence of the 23-kD fragment of the D1 protein in dark-grown *2pac ftsh1-1* cells might be due to a structural difference between PSII reaction centers from this strain, relative to the wild type. Therefore, we purified PSII from both the *2pac ftsh1-1* strain and the wild type so they could be compared. To enable purification of PSII from *2pac ftsh1-1*, we introduced a genetically encoded HIS-tag on the lumenal side of the PsbH subunit of PSII by crossing the *2pac ftsh1-1* mutant to the H-HIS strain (Cullen et al., 2007). Seven full tetrads were obtained. Progeny were genotyped by PCR to determine *FTSH1* and *RBD1* alleles, and one progeny (T7A) which contained the *2pac* and *ftsh1-1* mutant alleles was selected for further analysis **(Supplemental Fig. 5A**). Because *psbH* is a chloroplast-encoded gene and chloroplast inheritance is uniparental from the mating type + parent, all progeny were expected to express the HIS-tagged PsbH protein. Both the presence of PsbH-HIS and the absence of RBD1 were confirmed via immunoblot (**Supplemental Fig. 5B**).

T7A and H-HIS were grown on acetate in darkness, and the HIS-tagged PSII complex was purified by Ni-affinity chromatography. After isolation, the PSII samples were examined via BN-PAGE, and PSII monomers, dimers, and supercomplexes were observed in both H-HIS and T7A. (**Fig. 6A**). As shown in **Fig. 6B**, the PSII monomer from both H-HIS and T7A contained D1, D2, CP47, CP43, and HIS-tagged PsbH. To determine whether additional subunits might be lacking from the T7A dimer (or if assembly factors absent in H-HIS remained bound in T7A), the gel slices corresponding to the monomer were analyzed via LC-MS/MS (**Table 1**). The presence of D1, D2, CP47, CP43 and PsbH was confirmed as well as the additional presence of PsbO, PsbE, and PsbF in both H-HIS and T7A.

**Table 1:**
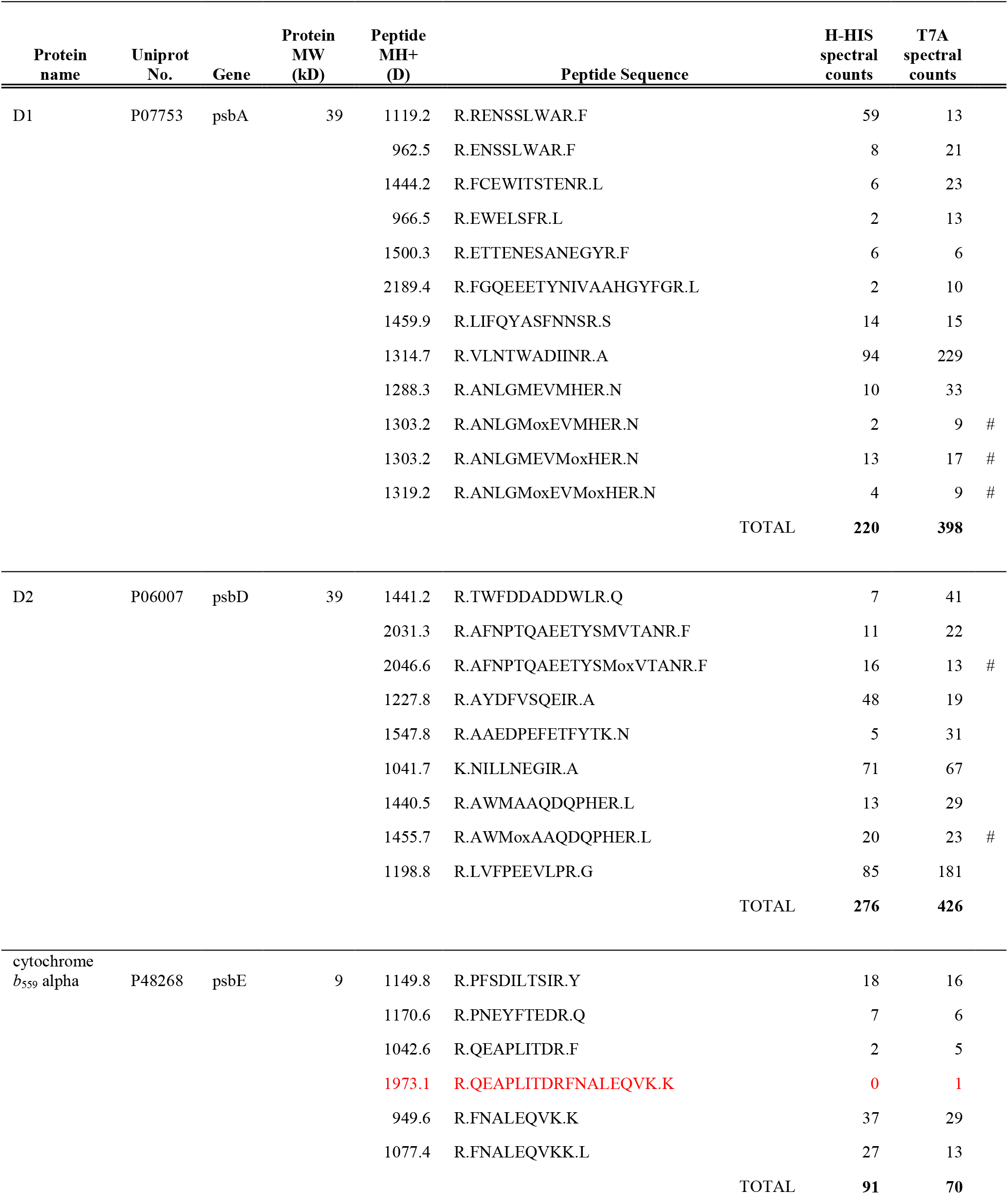
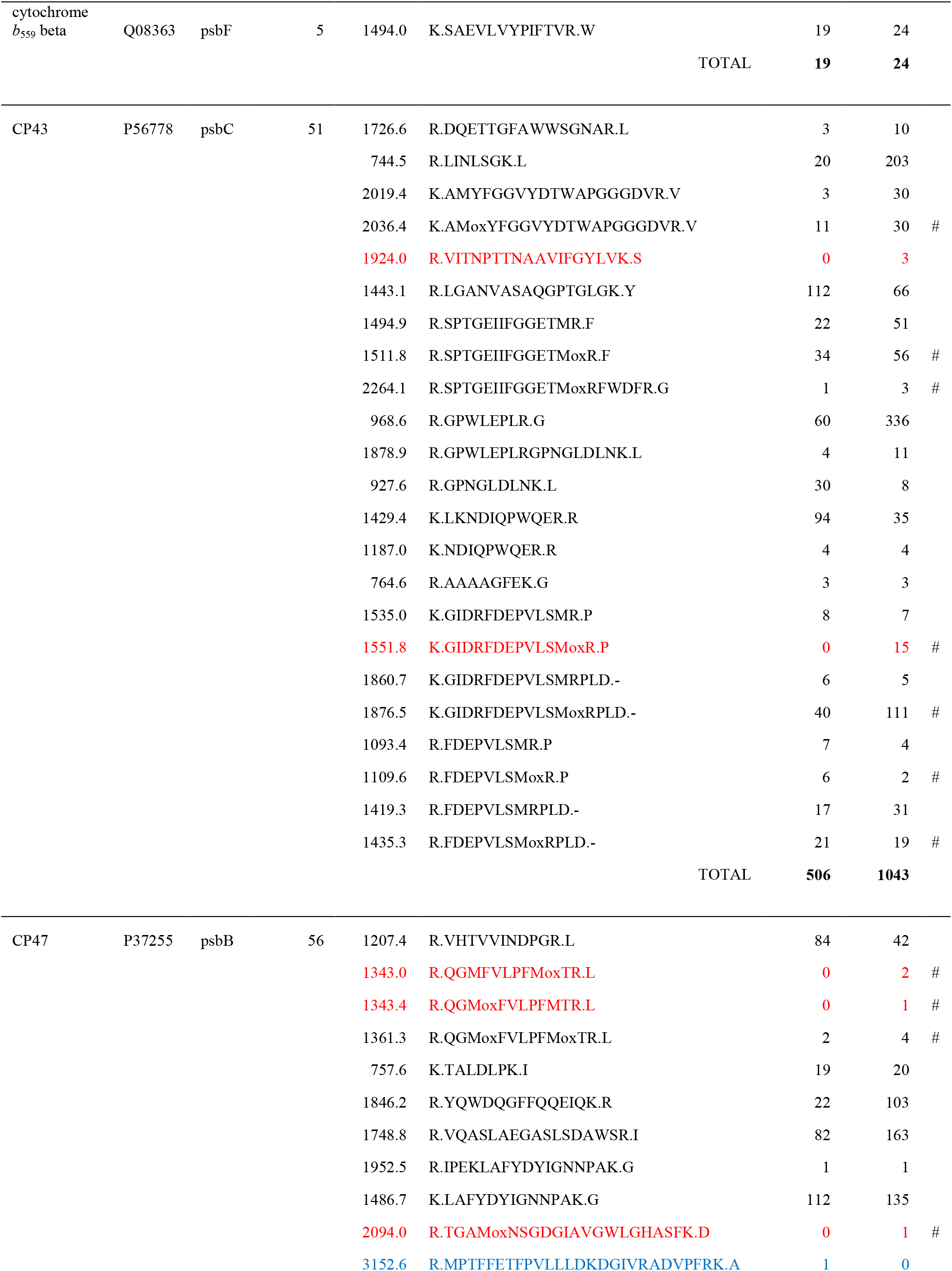
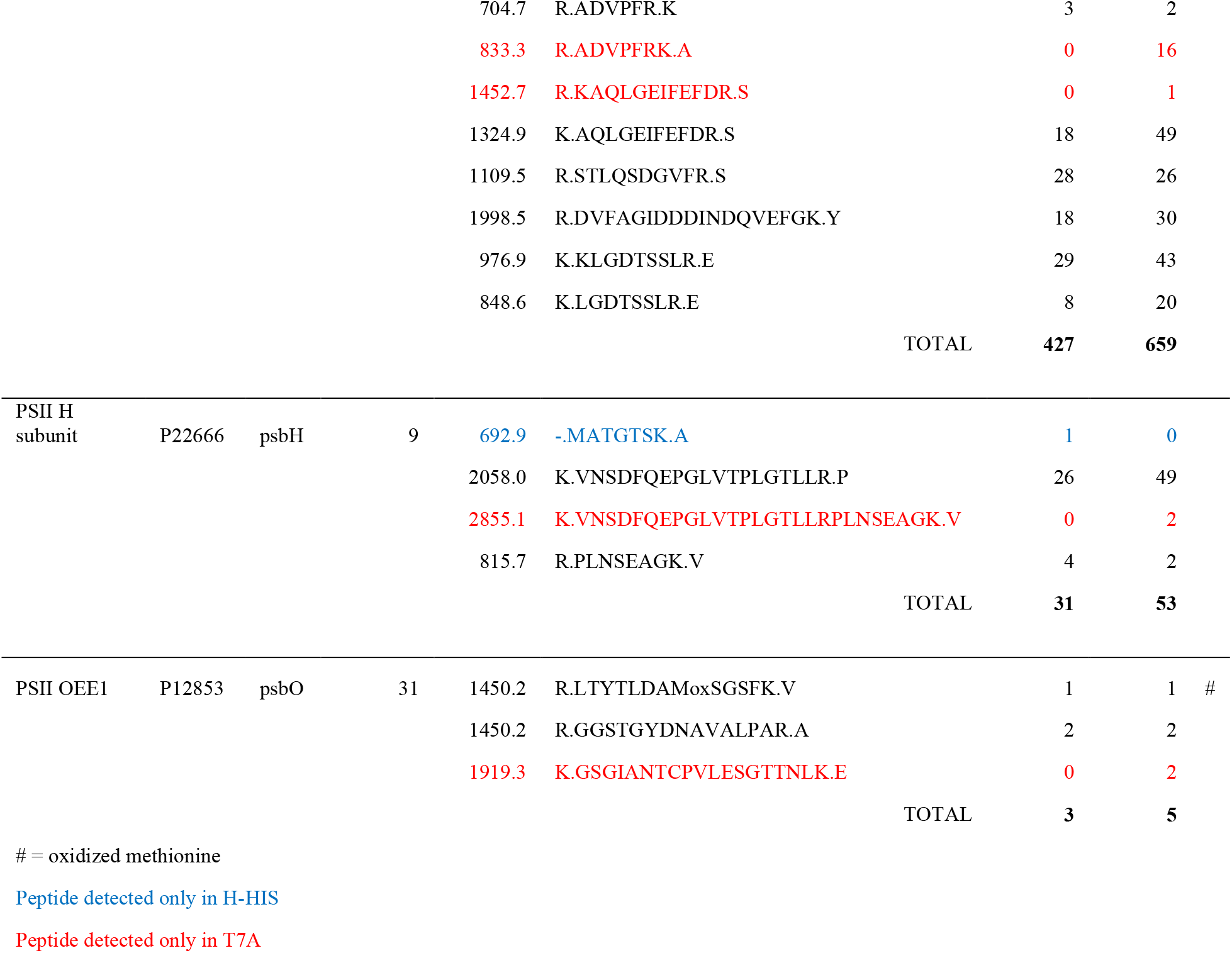
PSII proteins identified via LC-MS/MS analysis of isolated PSII dimers from H-HIS and H-HIS *2pac ftsh1-1* (T7A)

**Figure 6:**
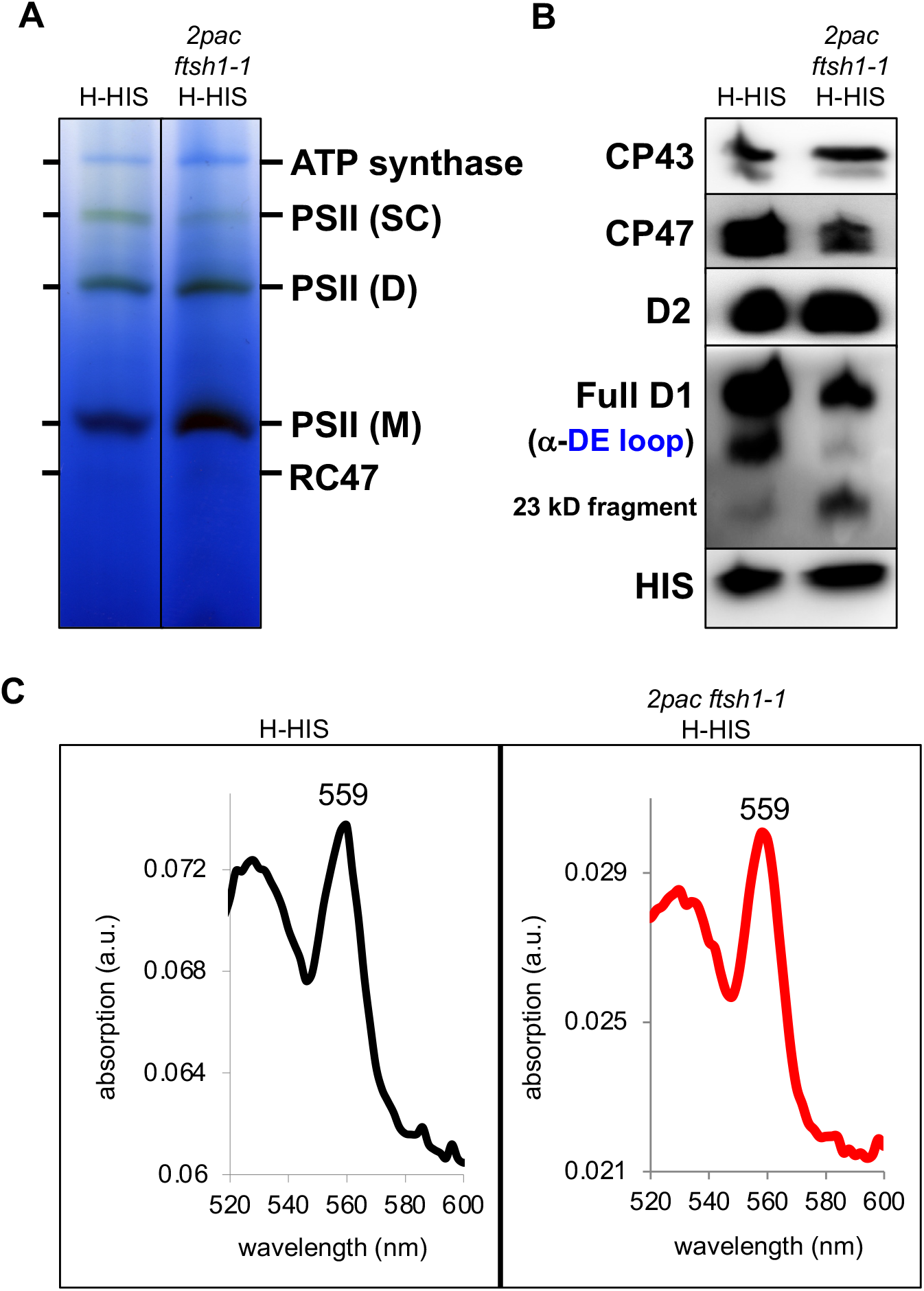
BN-PAGE, immunoblot analysis, and absorption spectroscopy of PSII purified from wild-type and *2pac ftsh1-1* backgrounds. **A)** BN-PAGE analysis of Ni-affinity-purified PSII from H-HIS (strain bearing a His-tagged PsbH subunit of PSII) and *2pac ftsh1-1* H-HIS (H-HIS strain with *2pac* and *ftsh1-1* mutations introduced via crossing) showing the accumulation of PSII supercomplexes (SC), dimers (D), monomers (M), and RC47 subcomplexes (RC47). **B)** Immunoblot analysis of excised bands corresponding to PSII monomers from both strains, showing presence of PSII subunits. **C)** Reduced minus oxidized absorption spectra of isolated PSIIs from both strains showing peaks at 559 nm, indicating the presence of the *b*-heme coordinated by cytochrome *b*_559_.

Given a previously hypothesized role for RBD1 in facilitating the photoprotection of nascent PSII complexes via an interaction with cytochrome *b*_559_ (García-Cerdán et al., 2019), we sought to examine whether the properties of cytochrome *b*_559_ were altered in the absence of RBD1. Both subunits of cytochrome *b*_559_ (PsbE and PsbF) were detected in PSII from T7A via mass spectrometry (**Table 1**), so we also tested for the presence of the heme coordinated by these proteins by recording reduced minus oxidized UV-vis absorption spectra of isolated PSII from H-HIS and T7A. As shown in **Fig. 6C**, both samples exhibited a strong peak centered at 559 nm, indicating that the PsbE and PsbF subunits detected by mass spectrometry do indeed coordinate the heme cofactor of cytochrome *b*_559_.

## Discussion

We have previously shown that the *Chlamydomonas 2pac* mutant lacking the *RBD1* gene is specifically deficient in PSII accumulation (Calderon et al., 2013). Our detection of wild-type levels of D1 and D2 protein synthesis in *2pac* rules out a role for RBD1 in enabling the translation of these PSII subunits. We did observe a modest reduction in CP47, CP43 and cytochrome *f* protein synthesis in *2pac* relative to WT, indicating that RBD1 may play a direct or indirect role in promoting their translation. However, the results from the immunochase experiment and the increased accumulation of PSII in the *2pac ftsh1-1* double mutant show that the low levels of PSII in *2pac* are due in major part to a post-translational instability of PSII subunits that can be partly alleviated by specific inactivation of the thylakoid-localized FtsH protease (**Fig. 1** and **Fig. 2**). These data are consistent with the results obtained on the cyanobacterial mutant lacking the RBD1 ortholog RubA (Kiss et al., 2019), in which the *ΔrubA* mutant was found to synthesize PSII subunits.

### PSII that accumulates in *2pac ftsh1-1* is highly sensitive to light and possibly structurally distinct from wild-type PSII

We found that *2pac ftsh1-1* strain was able to grow photoautotrophically (**Fig. 4A**) indicating that at least some amount of functional PSII was present since it was able to sustain growth. These results are consistent with the observation that the cyanobacterial Δ*rubA* mutant is able to grow photoautotrophically in the absence of RubA (Calderon et al., 2013; Kiss et al., 2019). We detected more PSII activity in *2pac ftsh1-1* when grown under low light on plates (**Fig. 4B**) than in strongly aerated cultures (**Fig. 4C**), suggesting oxygen- and/or light-dependent loss of PSII activity. The activity of PSII from *2pac ftsh1-1* was the highest in dark-grown cells, as evidenced by the partial recovery of variable fluorescence (**Fig. 5A**).

The D1 protein of PSII is the main target of photodamage (Ohad et al., 1984). The accumulation of D1 fragments in the single *ftsh1-1* is consistent with proposed models for plant (Kato et al., 2012; Kato and Sakamoto, 2018) and algal chloroplasts (Malnoë et al., 2014) in which there is a joint action between Deg proteases (endoproteolytic cuts) and FtsH proteases (processive degradation) during the repair of photodamaged PSII. Our detection of the 23-kD fragment of D1 in dark-grown *2pac ftsh1-1* cells (**Fig. 5B**) is highly unusual and suggests that the conformation of PSII in the mutant is altered. This fragment has been hypothesized to be either a degradation intermediate (De Las Rivas et al., 1992) or a product of translational pausing (Mullet et al., 1990). Reports presenting evidence that the fragment is due to degradation have hypothesized that it results from proteolysis (De Las Rivas et al., 1993; Keren et al., 1997; Spetea et al., 1999) or direct damage by reactive oxygen species (Miyao et al., 1995). In both cases, the trigger for degradation is thought to be photoinhibition, which would not occur in complete darkness. Indeed, the 23-kD fragment is not detected in dark-grown *ftsh1-1* (Malnoë et al., 2014). However, if the trigger were instead a conformational change occurring as a result of photodamage or misfolding (as might be the case in the absence of RBD1), the observed data would support a model in which RBD1 is required for the proper folding of D1 during PSII assembly (**Fig. 7**). A conformational change has indeed been previously suggested to act as a signal recognized by proteases that catalyze cleavage (De Las Rivas et al., 1993; Nakajima et al., 1996; Spetea et al., 1999). Specifically, occupation of the Q_B_ site by the PSII inhibitor PNO8 (*N*-octyl-3-nitro-2,4,6-trihydroxybenzamide) triggers the production of the 23-kD fragment in darkness, a process hypothesized to be due to PNO8-induced conformational changes at the DE loop (Nakajima et al., 1996). The stromal Arabidopsis Deg2 and Deg7 proteases have been shown to generate a 23-kDa and about 20-kDa fragment of D1, respectively, *in vitro* (Haussühl et al., 2001). Deg2 does not appear to have a role in D1 turnover after photoinhibition (Huesgen et al., 2006), but Deg7 participates in PSII repair in Arabidopsis *in vivo* (Sun et al., 2010). The *Chlamydomonas* genome encodes fourteen predicted Deg proteins (Schroda and de Vitry, 2022), 3 of which appear to be localized to the stroma. Based on sequence homology to two stroma-localized Arabidopsis proteins, Deg2 and Deg7 are likely found in the stroma (Schroda and Vallon, 2009; Schuhmann et al., 2012) while Deg1C, an ortholog of lumen-localized Arabidopsis Deg1, was experimentally identified in the stroma (Theis et al., 2019). Additionally, large-scale proteomic studies utilizing mass spectrometry have detected both Deg7 (Hemme et al., 2014) and Deg1C (Ramundo et al., 2014). The sole study dedicated to a *Chlamydomonas* Deg mutant is the characterization of the *deg1C* mutant which accumulates proteins involved in high light acclimation when grown at low light intensities (Theis et al., 2019). It is possible that Deg2, Deg7 and/or Deg1C function in PSII assembly quality control when RBD1 is absent, leading to production of the 23-kDa fragment in darkness and its accumulation in the *2pac ftsh1-1* strain but not in *ftsh1-1*.

**Figure 7:**
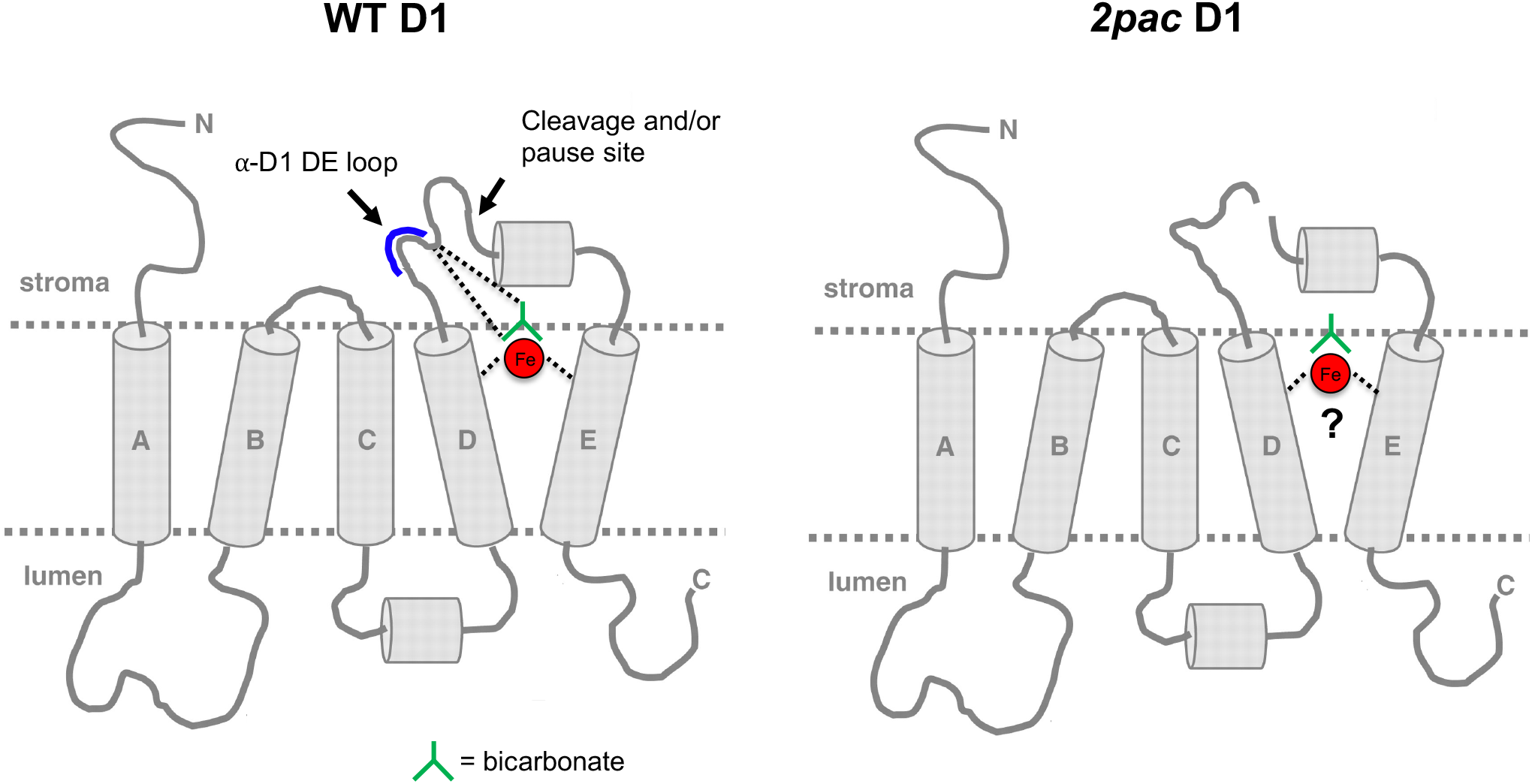
Model showing the effect of RBD1 mutation on the folding/maturation of D1. WT D1 (left side) provides four ligands that directly or indirectly (via bicarbonate, green) coordinate the non-heme iron of PSII (red sphere). These direct ligands are located in helix D (H215), helix E (H272), and in the DE-loop (E244 and Y246) close to the DE loop peptide (residues 234-242: NEGYRFGQE). A conformational change of the DE-loop of D1 may occur in the *2pac* mutant that lacks RBD1 (right side), resulting in proteolytic cleavage, possibly mediated by a Deg protease, followed by FtsH-mediated degradation. This conformational change may be due to the absence of the non-heme iron, which we propose may be delivered by RBD1 to D1 during the assembly of PSII (model adapted from (Hippler et al., 1998; Kapri-Pardes et al., 2007; Malnoë et al., 2014).

Alternatively, if the observed 23-kD fragment is a translational pause product (Mullet et al., 1990), then RBD1 might be required to facilitate the full translation of D1. This model, however, is not supported by the results of our pulse-labeling experiments in which we observed no differences in the rate of translation of full-length D1 in the *2pac* mutant (**Fig. 1A**), given our 7-minute pulse-labeling time which favors the detection of the rate of translation rather than the rate of degradation. However, the experiment requires a pulse under light since the incorporation of acetate is light-dependent, so it is still possible that pausing occurs in the dark.

The presence of aberrant PSII complexes in dark-grown *2pac ftsh1-1* would be consistent with the proposed role of RBD1 and its cyanobacterial ortholog RubA in PSII assembly (García-Cerdán et al., 2019; Kiss et al., 2019), but not with its separate proposed function in protecting assembly intermediates from photooxidative damage (García-Cerdán et al., 2019). The role in PSII assembly is further supported by the accumulation of RC47 and a smaller PSII assembly subcomplex detected via 2D-PAGE of *2pac ftsh1-1* thylakoids **(Fig. 3)** and in the observed interaction of RubA with two such PSII assembly subcomplexes during PSII assembly in cyanobacteria (Kiss et al., 2019).

### There are no detectable differences in protein composition of PSII from *2pac ftsh1-1* relative to the wild-type

The extreme light-sensitivity of PSII from *2pac ftsh1-1* compared to the wild-type led us to hypothesize that there might be a difference in protein and/or cofactor composition between the mutant and wild-type reaction centers. Mass spectrometry analysis of purified PSII complexes was unable to resolve any differences in the presence or absence of subunits (**Table 1**). This technique, however, suffers from the limitation that the detection of small hydrophobic proteins, of which there are many in PSII, is difficult. Consequently, we were unable to detect many of the small subunits of PSII, two of which (PsbJ and PsbL) are encoded by genes that lie directly downstream of *rubA* in the genomes of nearly all cyanobacteria (Calderon et al, 2013). It is therefore possible that one or more of these subunits is absent in the reaction centers purified from *2pac ftsh1-1*.

Our detection of cytochrome *b*_*559*_ in isolated PSII complexes indicates that the *b-*heme coordinated by PsbE and PsbF (cytochrome *b*_559_) is unaltered in the *2pac ftsh1-1* mutant and that RBD1 is unlikely to play a role in the maturation of cytochrome *b*_559_. While rubredoxins have generally been described as redox-active electron transport proteins (Lovenberg and Sobel, 1965), a recent report demonstrated that the rubredoxin fold of the enzyme 3-hydroxyanthranilate 3,4-dioxygenase (HAO) acts an iron reservoir (Liu et al., 2015). When iron is lost from the catalytic site of HAO, it is replenished by delivery of the so-called “spare tire” iron of the rubredoxin. RBD1 could conceivably serve a similar role for PSII, which binds a non-heme iron (**Fig. 7**). This non-heme iron is located at the electron acceptor side of PSII between the two quinones, Q_A_ and Q_B_. It is coordinated by four histidine residues, two from D1 and two from D2, and by a bicarbonate ion that provides a bidentate ligand (Umena et al., 2011; Kato et al., 2021). Many amino acid residues in the DE loop of the D1 protein, which are also in the vicinity of the non-heme iron (F239, Q241, E242, Y246), have been previously reported to undergo oxidation (Kumar et al., 2021), and the amino acids between 238 and 248 have been proposed to be the region where cleavage of the DE loop occurs *in vivo* (Greenberg et al., 1987). A model in which RBD1 functions in iron delivery is consistent with the observation that the cyanobacterial RubA interacts with D1 during the formation of the D1-D2 heterodimeric complex RCII (Kiss et al., 2019) but is not present in either of the direct precursors of RCII, the so-called D1_mod_ and D2_mod_ subcomplexes (Knoppová et al., 2022). The formation of the RCII complex is presumably accompanied by the insertion of the non-heme iron, given that both D1 and D2 provide the ligands for coordinating this metal (**Fig. 7**). Alternatively, RBD1/RubA could be involved in reduction of the non-heme iron (Calderon et al., 2013) during its ligation to the D1-D2 heterodimer. Both the non-heme iron of PSII and the iron-binding domain of RBD1/RubA are on the stromal side of the thylakoid membrane (Wastl et al., 2000; Umena et al., 2011; García-Cerdán et al., 2019; Kiss et al., 2019). Unfortunately, our attempts to detect the presence or absence of the non-heme iron of PSII via EPR and inductively coupled plasma-MS (ICP-MS) on isolated PSII from *2pac ftsh1-1* were unsuccessful. Nonetheless, the possibility that RBD1 delivers or reduces the non-heme iron during PSII assembly is compatible with all currently available data.

### The *ftsh1-1* mutant is a promising tool for studying mutations affecting thylakoid membrane complexes

Our work and a similar study that used the *ftsh1-1* mutation to accumulate and probe the function of a misassembled cytochrome *b*_6_*f* (Malnoë et al., 2011) have demonstrated the utility of this mutant in helping to understand PSII and cytochrome *b*_6_*f* biogenesis. Given the large number of FtsH-degradation targets in the plastid, future studies using the *ftsh1-1* mutant should provide insights into the biogenesis and function of a variety of other thylakoid membrane proteins.

## Materials and Methods

### *Chlamydomonas* strains, mutant generation, and growth conditions

Wild-type (4A+) and mutant strains *2pac* (Calderon et al., 2013), Fud7 (Bennoun et al., 1986), *ftsh1-1* (Malnoë et al., 2014), H-HIS (Cullen et al., 2007), and *2pac ftsh1-1* were grown at 25°C on Tris acetate-phosphate (TAP) medium (Gorman and Levine, 1965) or high salt (HS) minimal medium (Sueoka, 1960) as indicated. Strains were grown at various light intensities from dark to 30 μmol photons m^−2^ s^−1^, as indicated in the figure legends. Mutants were genotyped based on resistance to paromomycin (for *2pac*) or by PCR (for *2pac* or *ftsh1-1*) as described (Calderon et al., 2013; Malnoë et al., 2014) or via immunodetection with an anti-His antibody (H-HIS).

### Generation of *2pac ftsh1-1* double mutant

The *2pac* (mating type +) and *ftsh1-1* (mating type −) mutants were crossed. Four full tetrads, nine triads, and one dyad of the resulting progeny were assayed for the presence/absence of the *ftsh1-1* mutation by PCR (**Supplemental Fig. 1A**). To ensure the progeny were derived from successful mating, progeny were genotyped at the mating-type locus via PCR. All progeny groups showed a mixture of + and – mating types, indicating that they were derived from the cross (**Supplemental Fig. 1B**). Progeny were grown on TAP plates containing paromomycin to select for strains bearing the *2pac* mutation (**Supplemental Fig 1C**). One particular strain, T3.4, that contained both mutations was selected for further characterization and is the strain identified as *2pac ftsh1-1* when only one strain is shown and not specified.

### [^14^C]-acetate pulse-labeling

Exponentially growing cells at 2 × 10^6^ cells mL^−1^ from a 200 mL culture grown in TAP medium at 20 μmol photons m^−2^ s^−1^ were harvested by centrifugation, washed with minimum MIN-Tris medium and resuspended in 5 mL MIN-Tris medium at 2 × 10^7^ cell/mL. Cells were allowed to deplete their intracellular carbon pool for 1 hour under 20 μmol photons m^−2^ s^−1^ and strongly agitated for a good aeration. Afterwards, both the cytosolic translation inhibitor cycloheximide (final concentration 10 μg/mL) and [^14^C]-acetate (final concentration 10 μCi mL^−1^) were added simultaneously. Cells were allowed to take up the radiolabeled acetate for 7 min at 20 μmol photons m^−2^ s^−1^. The pulse-labelling was stopped by adding 35 ml of ice-chilled TAP medium containing 50 mM non-radioactive acetate followed by centrifugation at 4°C. Cells were resuspended in ice-chilled HEPES washing buffer, centrifuged, and immediately resuspended in ice-cold 0.2 M dithiothreitol and 0.2 M Na_2_CO_3_, frozen in liquid nitrogen and kept at −80°C until analysis by SDS-PAGE. Whole-cell protein extracts were separated by SDS-PAGE on a 12%-18% polyacrylamide gel in the presence of 8 M urea and visualized by autography. The assignment of the bands is based on mutant analysis (de Vitry et al., 1989; Kuras and Wollman, 1994)

### Immunochase

Inhibitors of chloroplast gene translation (chloramphenicol, 100 μg mL^−1^ and lincomycin, 500 μg mL^−1^) were added to 400 mL of exponentially growing cells in low light and TAP. Cells were incubated, with shaking, at 6 μmol photons m^−2^ s^−1^ over the course of 4 h, and 50 mL of cells were harvested at 0, 1, and 4 h post-addition of inhibitors before extraction of proteins, as described (Calderon et al., 2013). Protein samples were separated by SDS-PAGE on 12-18% polyacrylamide gels containing 8 M urea before transfer via semi-dry transfer system and subsequent blotting with specific polyclonal antibodies against ATP synthase (CF_1_ α and β subunits, PSII reaction center subunits D1-DE loop (Agrisera AS10704) and D2 (Agrisera AS06146), PSII core antennae CP43 and PSII extrinsic subunit OEE3.

### Chlorophyll fluorescence measurements

Chlorophyll fluorescence kinetics were measured at room temperature on dark-adapted cells. A home-built fluorimeter with a green detecting light was used for measurements on 1-mL aliquots of liquid cultures (Rappaport et al., 2007) before and after the addition of PSII-specific inhibitor 3-(3,4-dichlorophenyl)-1,1-dimethylurea (DCMU; 10 μM). A fluorescence imaging system (BeamBio, SpeedZen camera) with a blue detecting light was used for measurements of plates as described (Johnson et al., 2009). The maximum quantum yield of PSII photochemistry (F_v_/F_m_) was calculated as (F_m_-F_0_)/F_m_ where F_0_ is the fluorescence level of dark-adapted cells in the absence of DCMU F_m_ is the maximum level of fluorescence in the presence of DCMU or after a saturating light pulse.

### Analysis of thylakoid membrane proteins and complexes

SDS-PAGE and immunoblot analysis of proteins were performed as previously described (Calderon et al., 2013). Thylakoid membranes were isolated as described (Schottkowski et al., 2012). Briefly, cells were harvested at logarithmic growth phase (2 × 10^6^ cells mL^−1^) and washed in MKT buffer (10 mM Tricine-KOH, pH 7.5, 20 mM KCl, 25 mM MgCl_2_, 5 mM aminocaproic acid, 1 mM benzamidine and 0.2 mM PMSF) once before breaking by passage through a French pressure cell. Membranes were collected by centrifugation at 31,000 *g* for 30 minutes, then resuspended in ACA 750 (750 mM aminocaproic acid, 50 mM Bis-Tris, pH 7.0, 0.5 mM EDTA) to a concentration of 1 mg mL^−1^ chlorophyll. Membranes were solubilized by addition of an equal volume of ACA 750 containing 2% n-dodecyl β-D-maltoside (β-DM, Anatrace) for a final concentration of 0.5 mg mL^−1^ chlorophyll and 1% β-DM. Membranes were solubilized for 10 min on ice in the dark before centrifugation to pellet unsolubilized material. Solubilized membranes were then mixed 60:1 with loading buffer (100 mM BisTris-HCl, pH 7.0, 5% Coomassie G-250, 0.5 mM aminocaproic acid and 30% sucrose) and loaded onto a 4-16% precast BN-PAGE gel (Life Technologies). Second dimension analysis was performed by solubilizing BN-PAGE gel slices in 2x Laemmli buffer (Laemmli, 1970) and loading into precast 2D gels (Life Technologies).

Mass spectrometry experiments were performed at the Vincent J. Coates Protein Mass Spectrometry Facility. Bands were excised from BN-PAGE gels and subjected to analysis by reverse phase LC-MS/MS.

### Purification of HIS-tagged PSII

PSII purification was performed as described (Cullen et al., 2007), with the following minor modifications. Cultures were grown in 10 L bottles with vigorous stirring and bubbled with filtered air. All purification steps were performed in the dark, and all buffers were supplemented with betaine to a final concentration of 1 M. To account for differences in protein/chlorophyll content in the *2pac ftsh1-1* mutant, membranes from *2pac ftsh1-1* (T7A) were solubilized at final concentration of 0.6 mg mL^−1^ chlorophyll and 25 mM β-DM. Samples were further purified before analysis by MS by loading directly onto a precast 4-16% BN-PAGE gel (Life Technologies).

## Supplemental data

**Supplemental Figure 1: Genotyping of progeny from cross of *2pac* (mating type +) and *ftsh1-1* (mating type −) and isolation of double mutant *2pac ftsh1-1***.

**Supplemental Figure 2: Immunoblot analysis of progeny from *2pac ftsh1-1* cross showing steady-state levels of PSII subunits**.

**Supplemental Figure 3: *2pac ftsh1-1* strains display some variable fluorescence**.

**Supplemental Figure 4: Light-grown *2pac ftsh1-1* in liquid culture displays a very low PSII activity**.

**Supplemental Figure 5: Isolation of *2pac ftsh1-1* double mutants in the H-HIS background**.

## Funding

This work was supported by the Centre national de la Recherche Scientifique and Sorbonne Université (basic support to Unité Mixte de Recherche 7141), by the Agence Nationale de la Recherche (projects ANR-07-BLAN-0114 and ANR-12-BSV8-0011) and by the ‘Initiative d’Excellence’ program from the French State (Grant ‘DYNAMO’, ANR-11-LABX-0011-01).

## Acknowledgements

We thank Alizée Malnoë and Masakazu Iwai for extensive discussions, critical feedback, and experimental assistance. We thank Fabrice Rappaport, Alain Boussac, R. David Britt, Paul Oyala, David Marchiori, and Ruchira Chatterjee for technical assistance and helpful discussions. We also thank the *2pac* and *ftsh1-1* strains for inspiring two of their human caretakers to make a human genetic cross ten years after the creation of the algal genetic cross. This work used the Vincent J. Proteomics/Mass Spectrometry Laboratory at UC Berkeley, supported in part by NIH S10 Instrumentation Grant S10RR025622. K.K.N. is an investigator of the Howard Hughes Medical Institute. This article is subject to HHMI’s Open Access to Publications policy. HHMI lab heads have previously granted a nonexclusive CC BY 4.0 license to the public and a sublicensable license to HHMI in their research articles. Pursuant to those licenses, the author-accepted manuscript of this article can be made freely available under a CC BY 4.0 license immediately upon publication.

## Author Contributions

RHC, CdV, F-AW and KKN designed the research, analyzed the data and wrote the paper. RHC and CdV performed the experiments.

